# Striatal visual responses increase prior to visuomotor learning

**DOI:** 10.1101/2025.08.29.673104

**Authors:** Andrada-Maria Marica, Andrew J. Peters

## Abstract

The cortex and basal ganglia exhibit interdependent changes during learning. However, it is not clear whether plasticity occurs in sequence or concurrently across these structures. To address this question, we simultaneously recorded cortical and striatal activity while training mice on a visuomotor association task, which involved turning a wheel to move a stimulus from a cue location to a target location. Prior to the development of learned behavior, visual responses increased in the visual-recipient striatum. This was followed by the emergence of stimulus responses in both the medial prefrontal cortex (mPFC) and mPFC-recipient striatum from the onset of learned behavior. All of these regions also exhibited increased responses to stimuli in the rewarded target position, but while the visual-recipient striatum was non-selective between cue and target stimuli, the mPFC and mPFC-recipient striatum switched from being target-stimulus responsive before learning to being cue-stimulus responsive after learning. Our results suggest that sensorimotor learning involves routing stimulus information first to the sensory striatum, and then to frontal motor circuits.

## Introduction

Sensorimotor associations represent a fundamental type of adaptive behavior. These associations, which involve learning a behavioral response to an otherwise neutral stimulus, are thought to be formed by the cooperative action of the cortex and basal ganglia^1^. Supporting this idea, sensorimotor learning is impaired or eliminated when these structures are uncoupled via the corticostriatal pathway^2-4^. After learning, the sensory striatal domains become causally linked with the associated movement, such that behavior can be induced^5-8^ or inhibited^9-11^ through manipulation of either the striatum itself or corticostriatal input^12^. However, because the cortex and basal ganglia are connected in a loop, it is unclear whether sensorimotor learning involves the cortex “teaching” the basal ganglia, or if the basal ganglia “teaches” the cortex.

On one hand, the striatum generally inherits activity from the cortex through the corticostriatal pathway^13,14^. Learning is known to potentiate this connection^15-17^, yielding a more robust propagation of activity from the cortex to the striatum^13,18,19^. Striatal activity can then flow along subcortical routes to downstream motor regions^20,21^, providing a route for the striatum to drive behavior without further cortical involvement. In some cases, this may even culminate in the cortex only being involved during the learning phase, and being dispensable in favor of the striatum after learning^2,22,23^.

On the other hand, there is a prominent pathway from basal ganglia through the thalamus back to the cortex, in both closed- and open-loop configurations^5,24,25^. Striatal activity can therefore influence cortical activity^26-29^, which serves to reinforce cortical activity patterns^30-32^. In some cases, this makes the striatum only necessary for learning, and it is then dispensable for the execution of a learned sensorimotor association^33^.

One clue to deciphering the directionality of plasticity could be found in the temporal order of changes, where the cortex could lead, follow, or lag the striatum. We have previously observed specific changes in both of these structures during visuomotor learning, where stimulus responses increase in the visual-recipient striatum^13^, and a new stimulus response emerges in the medial prefrontal cortex (mPFC)^34^. Previous work has suggested that striatal changes may precede cortical changes^35-37^, and we therefore hypothesized that learning would be accompanied first by increased striatal responses, followed by the emergence of stimulus responses in the mPFC.

We tested this idea by recording cortical and striatal activity across sensorimotor learning, and indeed found that striatal changes preceded cortical changes. We recorded cortical and striatal activity simultaneously, which allowed us to determine the relative time course of changes, and also provided a means to precisely define striatal domains by their functionally correlated cortical regions. This revealed that striatal stimulus responses increased within the sensory cortex-recipient striatum prior to the development of associative behavior, while both the mPFC and mPFC-recipient striatum developed stimulus responses together and in step with behavioral changes. These results indicate that sensorimotor associations are preceded by strengthened sensory input into the striatum, and arise concurrently with sensory responses in the mPFC and mPFC-recipient striatum.

## Results

### Striatal visual responses increase with visuomotor learning

We trained mice on a simple visuomotor association task^34^. Mice were head-fixed in the center of three screens and rested their forepaws on a wheel that could be turned leftwards or rightwards (**Fig. 1A**). To initiate a trial, mice had to hold the wheel still for a randomly selected duration from 0.5-2s. At the beginning of each trial, a static vertical grating was displayed on the right-hand screen, and the position of the stimulus was yoked to the wheel. Turning the wheel leftwards would bring the stimulus to the center, where the stimulus would disappear, and a sucrose reward would be delivered. Turning the wheel rightwards enough to move the stimulus off the right-hand screen would end the trial. Between trials, there was a fixed inter-trial interval of 4-7s, when wheel turns had no effect.

**Figure 1.**
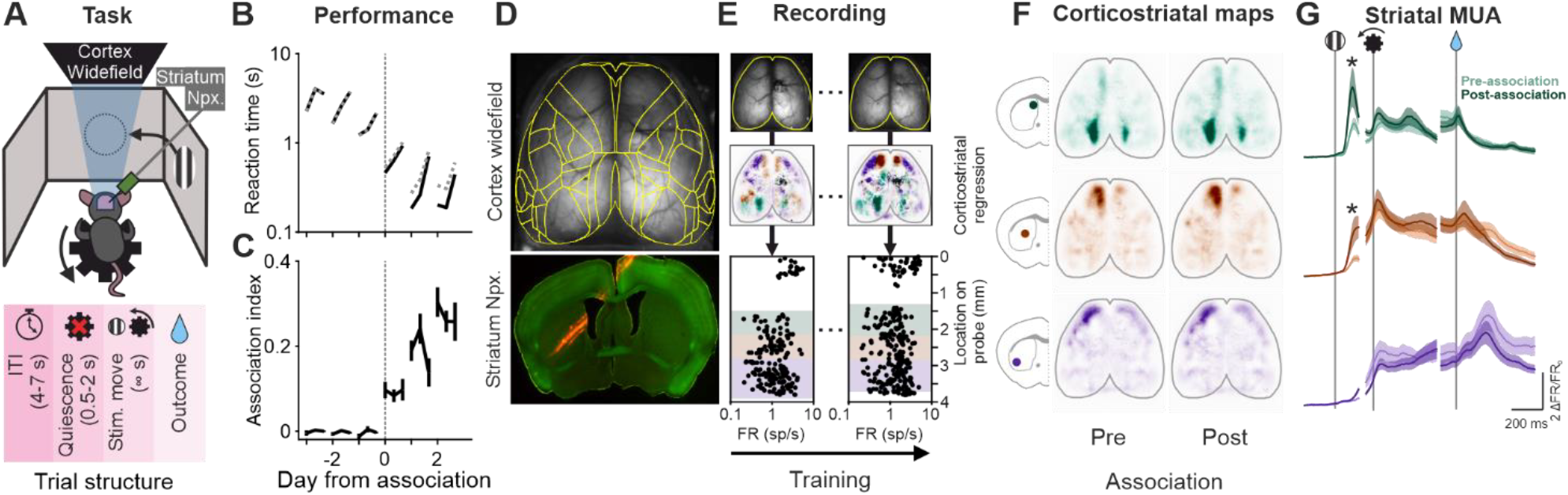
Visual- and mPFC-recipient striatal domains increase visual responses across visuomotor learning. (A) Top, schematic of behavioral setup; bottom, task trial structure. Npx., Neuropixels. (B) Mean reaction times measured (black) and by chance relative to stimulus (dotted gray). Sessions are split into thirds, aligned by association day, and averaged across mice (n = 14 mice). (C) Association index plotted as in (B), as the difference divided by the sum of the chance (gray) and measured (black) reaction times. Curves and error bars represent mean ± SEM. There is no change within the session before the association day (1-way ANOVA on day -1, p = 0.15). (D) Top. example average widefield image with CCF regions overlaid in yellow; bottom, example histology showing probe tract in red (E) Average widefield images (top), corticostriatal maps (middle), and electrophysiologically recorded units by probe location (bottom) for one animal across two recording sessions. Corticostriatal maps and units are colored by striatal domain. (F) Corticostriatal maps obtained from corticostriatal regression before and after learning, averaged within each domain and across animals. Corticostriatal maps do not change across learning (mean correlation of thresholded domain maps across learning vs. learning-shuffled within animal, p = 0.65 top, p = 0.16 middle, p = 0.26 bottom). (G) Time course of medium spiny neuron multiunit striatal spiking activity before (lighter colors) and after (darker colors) learning within each striatal domain. Activity is aligned to stimulus onset (left column), movement onset (middle column), and reward onset (right column). Stimulus-aligned activity increases in the visual and mPFC-recipient striatal domains (2-way ANOVA time x learning interaction, all three domains p < 1x10^-4^), while there is no change in movement-aligned and reward-aligned activity across domains (2-way ANOVA time x learning interaction, all three domains p = 1).

Mice learned an association between the visual stimulus and leftward wheel movement within a week. Wheel turns after stimulus onsets were initially rare, since mice were required to hold the wheel still before the stimulus was presented (**Fig. S1A, top**). Over training, the total number of wheel turns per session increased, and turns were consistently initiated after stimulus onsets (**Fig. S1A, bottom**). To determine whether mice were turning the wheel specifically in response to the stimulus, we measured reaction times as the duration from stimulus onset to movement onset, and tested whether these reaction times were shorter than chance using a conditional randomization statistic^34,38^. This method identified the first session in which each mouse began demonstrating visuomotor associative behavior, which was most often training day 4 (**Fig. S1C**). We could then align learning across mice relative to the first day that each animal exhibited associative behavior. For example, if a mouse learned the visuomotor association on training day 4, that day was defined as association day 0, while activity on training day 3 was defined as association day -1.

Visuomotor behavior developed across sessions rather than within a session. After determining the first day of associative behavior, we examined whether this association gradually appeared within a session, or whether it emerged from the start of a given session. We calculated the average reaction time of each session split into thirds, and aligned sessions relative to the first association day for each given mouse. This revealed a decreasing trend for reaction times across days, but an increasing trend within days, consistent with mice moving more slowly as they became sated or tired throughout each session (**Fig. 1B**). Both of these trends were present in the measured and chance reaction times, indicating that these effects were at least partially explained by changes in propensity to move the wheel that were not related to the stimulus (**Fig. 1B, black vs. gray lines**). We therefore quantified the degree of association as the difference between chance and measured reaction times divided by their sum, which provided an index of how much faster reaction times were compared to chance. As observed previously^34^, this demonstrated across-day increases in associative behavior after learning, rather than gradual increases within days (**Fig. 1C**).

A behavioral switch after learning was also observed in the efficiency of movements^33^. When examining the probability that any given wheel turn was preceded by a stimulus, stimuli and movements became less coincident at the end of sessions when the association had not been learned, while wheel turns became more efficiently executed in response to stimuli after learning (**Fig. S1D**). Together, the reaction times and movement efficiency indicate that visuomotor associative behavior emerged from the start of an identifiable day within each mouse.

We recorded cortical and striatal activity throughout training, and we functionally mapped striatal domains according to their cortical input regions. Cortical activity was measured across the dorsal cortical surface using widefield calcium imaging, and striatal spiking activity was recorded simultaneously using Neuropixels probes acutely inserted into the striatum on each session (**Fig. 1D**). The striatal probes were inserted along a diagonal trajectory spanning a functionally diverse swath from the dorsomedial to dorsolateral striatum (**Fig. 1D bottom, Fig. S2**). In order to precisely define the striatal domain being recorded by each part of the probe, we used linear regression to predict localized striatal multiunit activity from cortical fluorescence^13^. The kernels produced by this regression were maps of cortical regions that best correlated with striatal multiunit activity on a given segment of the probe (**Fig. S3A-B**). Across our recordings, we consistently recorded from three identifiable striatal domains, which were correlated with secondary visual cortex, medial prefrontal cortex (mPFC), and orofacial somatomotor cortex (**Fig. S3C**). We assigned each segment of each striatal recording to one of these domains based on their cortical maps, and then combined spikes from putative medium spiny neurons within each domain to produce domain-specific multiunit activity (**Fig. 1E, Fig. S3D-F**). This approach provides more interpretable results by contextualizing striatal activity within corticostriatal domains, and also accounts for trajectory variability both within and across mice.

Striatal visual responses increased across visuomotor learning, while corticostriatal topography remained stable. Cortical maps associated with each striatal domain likely correspond to upstream input regions, as previous results have demonstrated that they match anatomical projections and have a causal relationship to striatal activity^13^. Consistent with this anatomy being a fixed property, the cortical maps associated with each domain remained unchanged across learning (**Fig. 1F**). Each striatal domain exhibited activity related to its upstream cortical region, with visual stimulus-aligned activity in the visual-recipient striatum, movement-aligned activity in the mPFC-recipient striatum, and reward-aligned activity in the orofacial-recipient striatum (**Fig. 1G**). While movement- and reward-aligned activity did not change throughout the striatum, stimulus-aligned activity increased in both the visual- and mPFC-recipient striatum after learning (**Fig. 1G top and middle**). Visuomotor association learning therefore increases visual responses in the striatum without altering the functional mapping between cortex and striatum.

### Increased striatal visual responses precede learning

We examined the time course of activity in connected cortical and striatal regions relative to associative learning. Having observed an increase in striatal visual responses - across learning, we sought to characterize when these changes occurred relative to the emergence of associative behavior. In order to compare activity between connected cortical and striatal regions, we created cortical regions-of-interest from the corticostriatal regression maps (**Fig. S3C, yellow outlines**). This allowed us to directly relate multiunit spiking in a given striatal domain to fluorescence in the upstream cortical area.

Visual responses in the task increased in the visual-recipient striatum, and not in the visual cortex, before learning. The visual cortex exhibited both stimulus-aligned and movement-aligned responses throughout training (**Fig. 2A, top**). The movement response was likely due, at least in part, to the movement of the visual stimulus which was yoked to wheel movement. In contrast, the visual-recipient striatum exhibited only a weak stimulus response early in learning, while stimulus-aligned activity became robust in the couple days before learning (**Fig. 2A, bottom**). We compared stimulus-aligned activity after removing timepoints containing movement, and found that while stimulus responses did not change in the visual cortex, they increased in the downstream visual-recipient striatum (**Fig. 2B**). Importantly, this change preceded associative behavior (**Fig. 2B, black vs. dark gray lines**). The visual-recipient striatum therefore increased visual responses before learning, with no accompanying changes in the upstream visual cortex.

**Figure 2.**
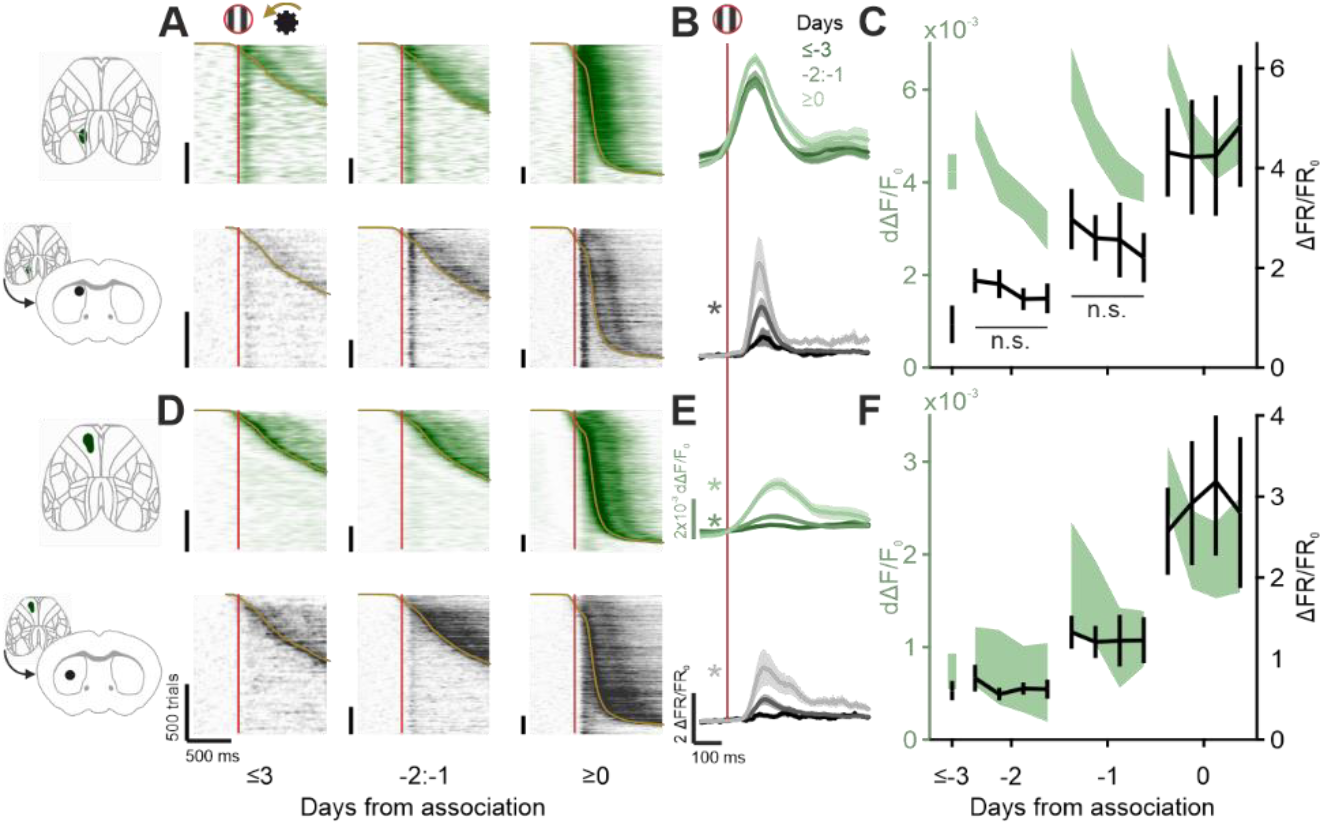
Increased visual responses in the task precede associative learning. (A) Activity across all trials from all mice in the visual cortex (top, green) and visual-recipient striatum (bottom, black), sorted by reaction time and grouped by day from association. Red lines are stimulus onset, yellow lines are movement onset. Heatmaps are vertically smoothed with a moving mean of 20 trials to display trends across trials. (B) Time course of average stimulus-aligned activity from the regions in (A), excluding timepoints following 100ms before movement onset for each trial to exclude movement-related activity. Colors are days from association, curves and shading are mean ± SEM across mice. The visual-recipient striatum has an increased stimulus-aligned response before learning (difference in maximum activity 0-0.2 s after stimulus onset vs. shuffled day group within mouse, days ≤ -3 vs -2:-1 p = 0.004), while there is no change in the visual cortex (same statistic, p = 0.46). (C) Stimulus-aligned activity within sessions. Sessions are split into quartiles, trials with reaction times >300ms are averaged together, and the maximum of that average within 200ms of stimulus onset is plotted. Curves and error bars or shading are mean ± SEM across mice. Activity does not increase within any day in the visual-recipient striatum (1-way ANOVA, day -2 p = 0.67, day -1 p = 0.84, day 0 p = 0.98), while activity decreases within each day in the visual cortex (1-way ANOVA, day -2 p = 9.2x10^-4^, day 1 p = 0.002, day 0 p = 0.004). (D-F) Data as in A-C, for the mPFC domain (top, green) and mPFC-recipient striatum (bottom, black). For (E), there is an increase in stimulus-aligned activity both before and after the association in the mPFC (statistic as in (B), days ≤ -3 vs -2:-1 p = 0.005, days -2:-1 vs ≥ 0 p < 1x10^-4^), and an increase mPFC-recipient striatum after learning (same statistic, days ≤ -3 vs -2:-1 p = 0.21, days -2:-1 vs ≥ 0 p = 0.001).

The visual-recipient striatum increased visual responses between days, rather than within a day. Visual responses could have increased gradually within a training session, or otherwise could have increased between days. We distinguished these possibilities by splitting each session into quartiles, removing trials with reaction times less than 300 ms to exclude movement, and quantifying peri-stimulus activity within each corresponding group of trials. In the visual-recipient striatum, visual responses did not increase within training sessions, with increases instead only occurring between the end of a previous session and the start of the next session (**Fig 2C, black lines**). As noted above, visual responses in the visual cortex did not increase across sessions, although there was a decreased response within each session possibly as an effect of decreasing arousal or task engagement (**Fig. 2C, green shading**). Visual responses in the visual-recipient striatum therefore increased between days prior to learning.

The mPFC also exhibited increases in task visual responses before learning. Unlike the visual cortex and visual-recipient striatum, the mPFC and mPFC-recipient striatum were completely unresponsive to visual stimuli early in learning, although they exhibited movement-aligned activity (**Fig. 2D**). The mPFC then developed a small stimulus response in the couple days before learning (**Fig. 2D-E, top row**). This response increased further after learning, and it was accompanied by an increased stimulus response in the mPFC-recipient striatum (**Fig. 2D-E**). These increased visual responses also emerged between days, rather than increasing gradually within days (**Fig. 2F**). The mPFC therefore increased visual responses prior to learning, although this was not consistently observed in the mPFC-recipient striatum.

The orofacial cortex and orofacial-recipient striatum did not exhibit visual responses during the task. As expected from the orofacial regions, activity was aligned to movement and not stimulus onset, with no visual responses present throughout training (**Fig. S4A-C**). Visual responses in the striatum were therefore restricted to the visual- and mPFC-recipient domains.

### Stimulus context-dependence and selectivity differs by striatal domain

In addition to examining visual responses in the context of the task, we also presented visual stimuli in a passive context. After each training session, stimuli at three different positions were passively presented fifty times each, during which time turning the wheel had no effect. Mice rarely turned the wheel during these passive presentations, and any trials with wheel movement were excluded from analysis. These passive presentations served three purposes. First, we could determine whether visual responses were consistent across contexts, or were instead modulated by task engagement. Second, we could present various stimuli to test the selectivity of changes in visual responses. Third, visual activity could be examined in the absence of movement, which was not possible in the task after learning when stimulus onsets and movements were not well separated.

Passive responses to the task cue stimulus increased before learning in the visual-recipient striatum. We first examined responses to stimuli in the cue position on the right-hand screen, which were identical to stimuli presented at the start of each trial during the task (**Fig. 3A**). The visual cortex was consistently responsive to this stimulus across training, while responses increased in the visual-recipient striatum in the couple days prior to associative behavior (**Fig. 3B**). This change mirrored what was observed during the task, and reinforced the finding that increased visual responses in the visual-recipient striatum precede learning and are not related to changes in the upstream visual cortex.

**Figure 3.**
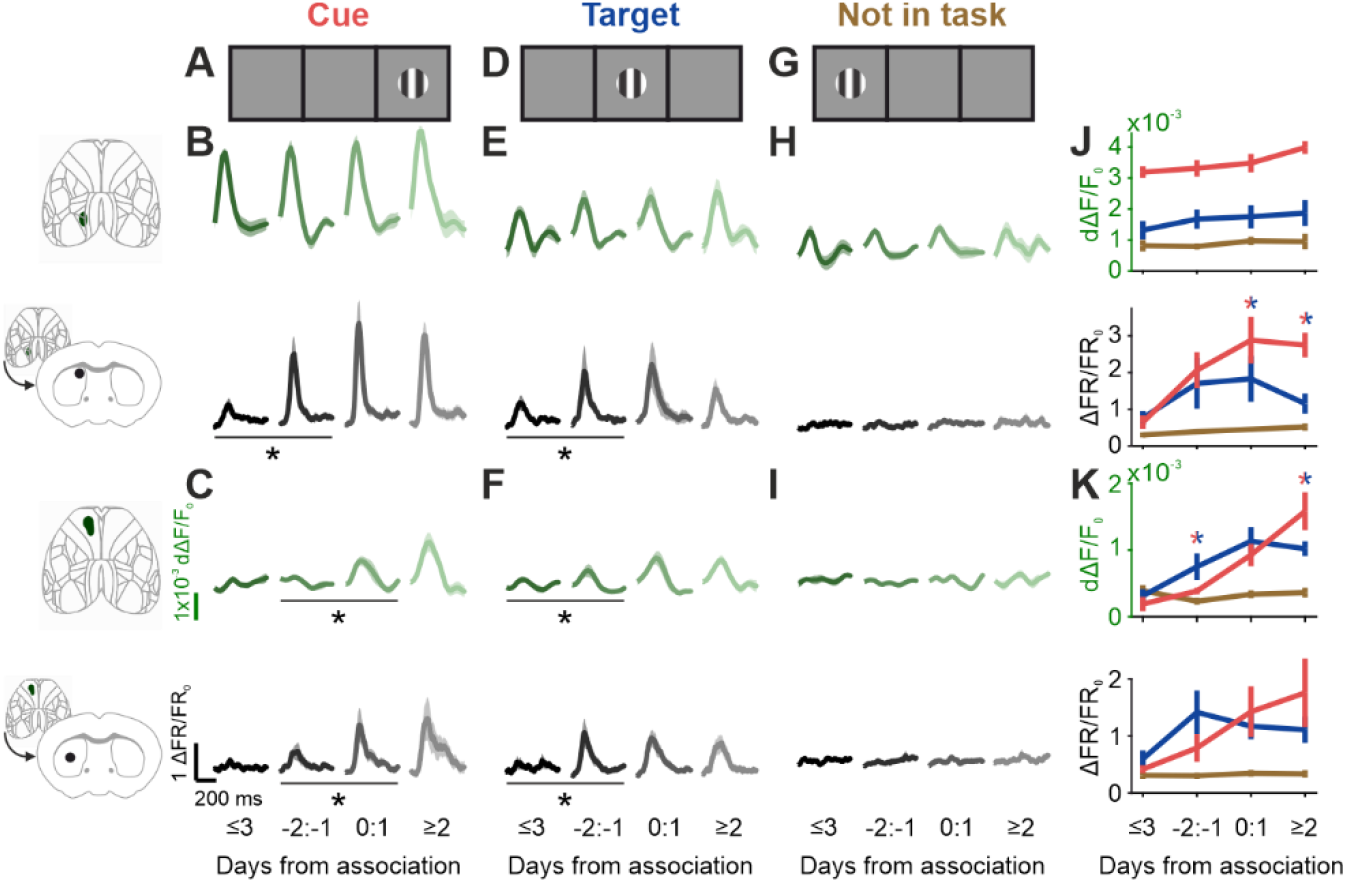
Passive visual responses to task-relevant stimuli increase in the visual-recipient striatum before associative learning. (A) Schematic of stimulus positioned at the task cue location on the right-hand screen. (B) Time course of activity aligned to the onset of the task cue position stimuli in the visual cortex (top, green) and visual-recipient striatum (bottom, black). Curves and shading are mean ± SEM across mice. Activity increases in the visual-recipient striatum before learning (difference in maximum activity 0-0.2 s after stimulus onset vs. shuffled day group within mouse, days ≤ -3 vs -2:-1 p = 0.0038), and does not change in the visual cortex (same statistic, p = 0.17). (C) Data as in (B) but for the mPFC and the mPFC-recipient striatum. Activity does not change before learning (statistic as in (A), days ≤ -3 vs -2:-1 mPFC p = 0.1, mPFC-recipient striatum p = 0.13), and increases after learning both the mPFC (same statistic, days -2:-1 vs ≥ 0 p = 0.001) and mPFC-recipient striatum (same statistic, p = 0.034). (D-F) As in (A-C), for stimuli positioned in the task target position on the center screen. Activity increases before learning in the visual-recipient striatum (statistic as in (A), days ≤ -3 vs -2:-1 p = 0.026), mPFC (same statistic, p = 0.018), and mPFC-recipient striatum (same statistic, p = 0.012). (G-I) As in (A-C), for stimuli in a position not shown during the task on the left-hand screen. Activity does not change in any region (statistic as in (A), days -2:-1 vs ≥ 0 visual cortex p = 0.14, visual-recipient striatum p = 0.15, mPFC p = 0.06, mPFC-recipient striatum p = 0.28). (J) Maximum amplitude of average stimulus-aligned activity within 200 ms of stimulus onset, for visual cortex (top) and visual recipient striatum (bottom). Colors are stimulus position, curves and error bars are mean ± SEM across mice. Activity is higher for cue (right-hand) stimuli compared to target (center) stimuli in the visual-recipient striatum after learning (difference between activity for each stimulus vs. shuffled day group within mouse, days 0:1 p = 0.005, days ≥ 2 p = 0.008). (K) Data as in (J) for the mPFC and mPFC-recipient striatum. In the mPFC, activity is higher for task target stimuli than task cue stimuli before learning (statistic as in (J), days -2:-1 p = 0.05), while activity is higher for task cue stimuli than task target stimuli after learning (statistic as in (J), day ≥ 2 p = 0.03).

The mPFC and mPFC-recipient striatum did not exhibit changes in cue stimulus responses before learning. Although the mPFC showed weak stimulus activity during the task in the couple days prior to associative behavior, neither had consistent responses to the cue stimulus when it was passively presented. Instead, both the mPFC and mPFC-recipient striatum developed a response to the passive cue stimulus only from the first day of associative behavior (**Fig. 3C**). This is consistent with our previous findings that the mPFC becomes visually responsive synchronously with learning^34^, and suggests that the responses observed during the task before learning were sensitive to context.

Passive responses to stimuli at the task target location increased across all visually responsive areas before learning. In addition to presenting the stimulus at the cue position where it appeared at the onset of each trial, we also presented the stimulus at the target position on the center screen, which was the location that would trigger a sucrose reward in the task (**Fig. 3D**). As with the cue stimulus, the visual cortex was stably responsive to the target stimulus throughout training. However, the visual-recipient striatum, the mPFC, and the mPFC-recipient striatum all exhibited increased responses to the target stimulus in the couple days before learning (**Fig. 3E-F**). The cue and target stimulus responses were contrasted by a lack of response to stimuli on the left-hand screen (**Fig. 3G-I**), which was expected both because this stimulus was not shown during the task, and because it was ipsilateral to the recorded striatum and corresponding cortical regions of interest. The visual-recipient striatum therefore increased responses to both task-relevant stimuli before learning, while the mPFC and mPFC-recipient striatum increased responses to the target stimulus before learning, and the cue stimulus after learning.

The visual-recipient striatum and mPFC became selectively responsive to the cue stimulus compared to the target stimulus after learning. While the visual-recipient striatum increased passive responses to both the cue and target stimuli before learning, responses to the target stimulus became weaker than the cue stimulus after learning (**Fig. 3J**). A similar trend was observed in the mPFC (and non-significantly in the mPFC-recipient striatum), but while responses were larger for the target stimulus than the cue stimulus before learning, this selectivity reversed after learning (**Fig. 3K**). Together, this demonstrates that increased responses to the cue stimulus persist after learning, while target responses become weaker than cue responses after learning.

The orofacial cortex and orofacial-recipient striatum did not exhibit passive visual responses, consistent with their lack of a visual response during the task (**Fig. S4D-E**).

### Stimulus selectivity and preference differs by striatal cell type

Striatal neurons could be classified into putative cell types from their electrophysiological properties. The above results focused on the multiunit activity of putative medium spiny neurons (MSNs), though putative fast-spiking interneurons (FSIs) could be identified from their short waveforms, and tonically active neurons (TANs) as putative cholinergic interneurons could be identified from their wide waveforms and regular firing rate (**Fig. S5**).

Visually responsive neurons across all three cell types were selective to cue or target stimuli. Having observed responses to passively presented stimuli in both the task cue and target positions, we investigated whether individual neurons were selective or co-active to these stimuli. We first identified neurons which were significantly modulated by stimulus presentation at either location. Visually responsive neurons could be found across all three cell types, and cue-responsive neurons exhibited stimulus-aligned activity in both the passive and task contexts (**Fig. 4A**). MSNs, FSIs, and TANs were all predominantly selective to stimuli at either the cue or target position, and were less co-responsive than expected by chance (**Fig. 4B**).

**Figure 4.**
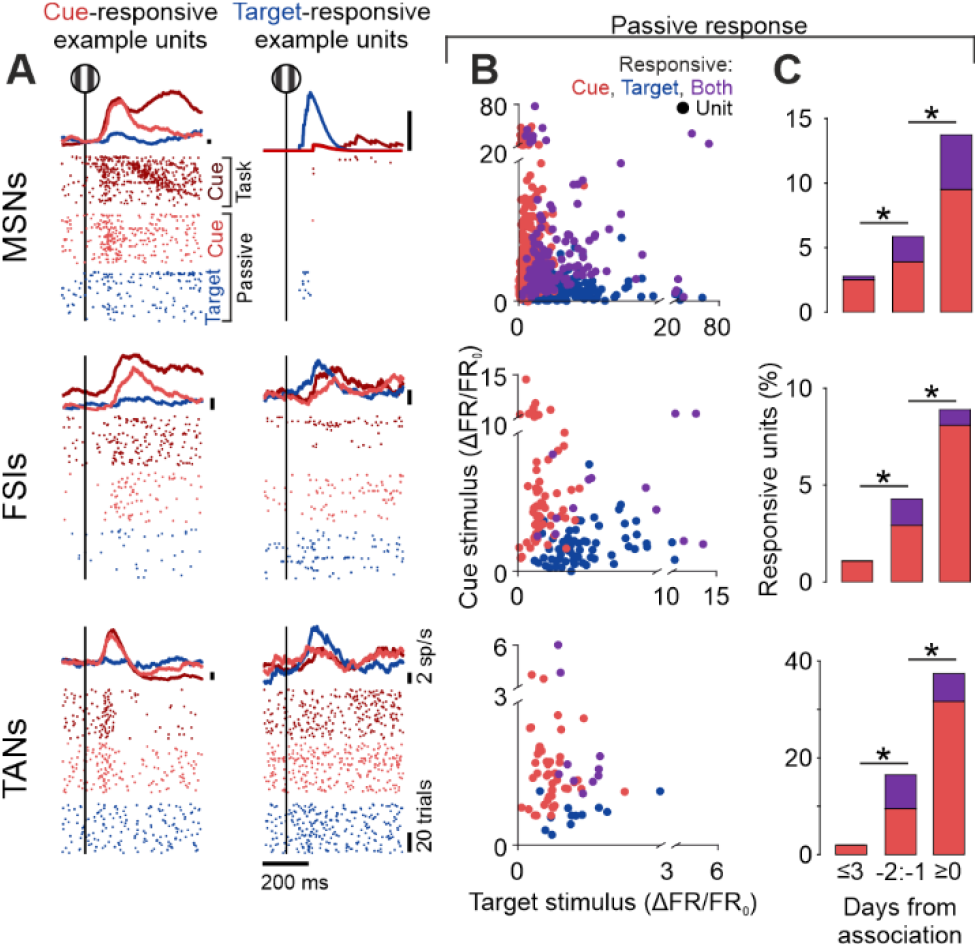
All striatal cell types are selective to stimuli and increase visual responses before learning. (A) Peri-stimulus time histograms (PSTHs) and raster plots of stimulus-aligned activity in the task and passive contexts for example neurons. Top, medium spiny neurons (MSNs); middle, fast-spiking interneurons (FSIs); bottom, tonically active neurons (TANs). Dark red is stimulus onset during task, light red is passive cue stimulus, blue is passive target stimulus. (B) Maximum activity within a 50-150ms window after stimulus onset, averaged across trials for each stimulus, and plotted as cue position response (y-axis) against target position response (x-axis). Each dot is a neuron that is responsive to the cue and/or target stimulus, with color denoting stimulus responsiveness. Only neurons within the visual- and mPFC-recipient striatum are included. Rows are cell type as in (A). In all three cell types, neurons responsive to both cue and target stimuli co-occur less frequently than expected by chance (fraction of co-responsive neurons vs. shuffled responsiveness label across units within animal, day group, and domain, MSN p = 1x10^-4^, FSI p = 1x10^-4^, TAN p = 0.034). (C) Fraction of units in the combined visual- and mPFC-recipient striatum responsive to stimuli in the cue position, either alone (red) or co-responsive to stimuli in the target position (purple). Rows are cell type as in (A). Units are grouped relative to day from association. Cue stimulus-responsive neurons increase in all cell types both before and after learning (fraction of cue-responsive neurons vs. shuffled day group within animal and domain, MSN days ≤ -3 vs 2:1 p = 0.003, days 2:1 vs ≥ 0 p < 1x10^-4^; FSI days ≤ -3 vs 2:1 p = 0.047, days 2:1 vs ≥ 0 p = 0.009; TAN days ≤ -3 vs 2:1 p = 0.0021, days 2:1 vs ≥ 0 p = 0.0011).

All cell types exhibited an increase in cue responses prior to learning. Consistent with the changes seen in multiunit activity, the fraction of responsive MSNs increased before mice exhibited associative behavior (**Fig. 4C, top**). The fraction of cue-responsive neurons also increased in both the FSI and TAN populations in the couple days preceding learning (**Fig. 4C, middle and bottom**). Therefore, even though the TANs receive their strongest input from different sources than the MSNs and FSIs^39^, all of these cell types increase their visual responses before a visuomotor association is learned. This pre-learning increase was followed by a further increase in cue-responsive neurons in all three populations after learning (**Fig. 4C**).

## Conclusion

Our results show that stimulus responses increase in the visual-recipient striatum prior to learning a visuomotor association. This suggests that robust striatal stimulus responses may be a necessary antecedent for linking those stimuli with actions.

Striatal stimulus responses are likely increased by corticostriatal potentiation^15^, as we observed no change in upstream visual cortical activity. This form of potentiation can include slow structural changes^16,17^ and may require sleep to fully develop^18^, consistent with our observations that changes occurred between days rather than within sessions.

The trigger for striatal potentiation may be coincident input from the visual cortex and dopaminergic projections^40^. In this case, it is unclear what key event would drive dopamine release in the dorsomedial striatum before learning. One possibility is that the visual stimulus itself drives dopamine responses, but while these have observed in some naive mice^41^, they are often completely absent^42^ and only consistently present in trained mice^43^. Furthermore, these dopaminergic stimulus responses have been suggested to emerge concurrently with learning, rather than preceding it^42,44^. It is therefore possible that stimulus-driven dopamine responses do not drive the initial learning, and only emerge later to enable modification of learned behavior^45,46^. Instead, dopamine responses in the naïve dorsomedial striatum have also been observed for both movement^47^ and reward^42,45^, albeit weakly. In our task, mice exhibit a non-specific increase in wheel movement before they learn the sensorimotor association, which would bring these movement- or reward-related dopamine signals within a critical time window from visual input by chance^48,49^.

Another component for striatal sensory potentiation in our task may be cholinergic signals from the TANs^50^. We found that TANs developed stimulus responses prior to associative behavior, mirroring the change in MSNs and FSIs. However, TANs receive proportionally more of their input from the thalamus^39^, suggesting that visual information into the striatum increases along both the corticostriatal and thalamostriatal pathways before sensorimotor learning. The source of thalamostriatal excitation is unclear, as the upstream thalamic nuclei can be driven either by the cortex or superior colliculus^51^, but our results may indicate the involvement of other subcortical structures in striatal potentiation^52^. The impact of TAN activation is also unclear, as TANs can both directly modulate MSNs^53^ and regulate dopamine release^54^.

Visual responses exhibited different changes depending on the striatal domain, suggesting a dissociation of roles by region. In the visual-recipient striatum, we found that responses to both cue-position and target-position stimuli increased before learning. This may indicate that the visual-recipient striatum acts as a gate for learning, where striatal responses first develop for all behaviorally relevant stimuli. Once responses to a given stimulus have been potentiated in the visual-recipient striatum, they can then likely serve to reinforce behavior in response to that stimulus. This principle of striatal activity reinforcing behavior has been demonstrated by artificially exciting the striatum, which has the effect of reinforcing any given ongoing movement^55-57^. Stimulating dopamine release in the striatum also has a similar effect, again emphasizing a role for the striatum in behavioral reinforcement^58,59^. The pathway by which behavior is reinforced may be through subcortical structures^20,21,60^. However, given our observations of subsequent changes in the mPFC, an interesting possibility is that striatal activity induces mPFC plasticity through an open-loop pathway back to cortex^25^.

In contrast to the visual-recipient striatum, the mPFC and mPFC-recipient striatum both exhibited responses to the target stimulus before learning, and responsiveness switched to the cue stimulus after learning. The simultaneous change between regions suggests that activity already propagates from the mPFC to the mPFC-recipient striatum without the prerequisite of corticostriatal strengthening. This naively robust pathway may underlie fast modification of a previously learned association^61,62^, which does not induce changes in sensory-recipient striatum^11,63,64^. The stimulus selectivity in these regions suggests that activity may be at least partly related to reward-predicting value rather than behavioral association^65^, as the target stimulus is more temporally locked to rewards early in learning and therefore may be perceived by the animal as more valuable. However, activity in the mPFC is causally involved in executing a learned visuomotor association^66^, and therefore may act in both sensorimotor and value-prediction roles.

Together, our results suggest that the direction of influence between the cortex and striatum may depend on the region or context. For example, the cortex-to-striatum route may be a key component of the sensory system, where the sensory corticostriatal pathway is weak in naïve animals and requires potentiation to enable behavioral learning^15^. Conversely, the striatum-to-cortex route may be a key component of the motor system, where cortical activity patterns are shaped by striatal influence^67^. Both of these directions may also be used simultaneously, which could shape the activity of both structures towards a common function^68^, or could develop separable functions carried out by each structure^69^.

## Data and code availability

The data used to generate the figures are available as downloadable files on OSF at https://osf.io/ynjdz/.

The code used to generate the figures is available on GitHub at https://github.com/PetersNeuroLab/PetersLab_papers/tree/main/%2BMarica_2025.

## Acknowledgements

We thank Da Song and Haron Avgana for helpful discussions. This work was supported by a Sir Henry Dale Fellowship jointly funded by the Wellcome Trust and the Royal Society (224156/Z/21/Z) to A.J.P.

## Author contributions

A.-M.M. and A.J.P. contributed to all aspects of this paper.

## Methods

### Animals

All experiments were conducted according to the UK Animals (Scientific Procedures) Act 1986 under personal and project licenses issued by the Home Office. Animals were adult (6 weeks or older) male and female transgenic mice (tetO-GCaMP6s; CaMKII-tTa^70^).

### Surgery

Mice underwent two surgeries each: one for headplate implantation and widefield imaging preparation, and a second one to perform a craniotomy for acute electrophysiology.

For the headplate implantation surgery, animals were anesthetized with isoflurane and their heads were shaved. They were then fitted into a stereotax (RWD), placed on a feedback-controlled heat pad (ThermoStar, RWD), and injected subcutaneously with meloxicam and buprenorphine. The scalp was cleaned with iodine and ethanol, and cut away to expose the skull. The periosteum was cut from the skull, and muscles were detached from the occipital bone. The skin was glued around the incision site using VetBond (3M), and a custom titanium headplate, consisting of a bar over the occipital bone and a U-shape surrounding the rest of the skull, was affixed to the occipital bone with dental cement (C&B-Metabond, Parkell). Clear dental cement was then was applied within the U-shape of the headplate to cover the skull and fix the headplate to the skull at the front and sides. UV-curing optical glue (NOA81 and Opticure LED, Norland) was applied on top of the clear dental cement to improve transparency. Mice were provided meloxicam-infused jelly after surgery, and given at least 5 days to recover.

The craniotomy surgery took place after rig habituation and naïve widefield recordings. The craniotomy was performed at approximately 0.6 mm AP and 0.7 mm ML relative to bregma. Mice were anesthetized and then received a subcutaneous injection of meloxicam. The optical glue and dental cement were drilled away over the area of the craniotomy. The craniotomy was performed using a 1.5-2mm biopsy punch, and in some cases a durotomy was performed. The exposed craniotomy was then covered with Kwik-Cast (WPI), and the Kwik-Cast was covered by a titanium lid screwed into the headplate. Recordings and training began the day after the craniotomy was performed.

### Habituation and naïve widefield recordings

After mice recovered from the headplate surgery, they were head-fixed in the rig briefly across three days. This acclimated mice to handling, head-fixation, and the wheel under their forelimbs. Mice were also given sporadic bursts of sucrose water drops during these sessions, which taught them about the presence of the sucrose reward, and also likely linked the sucrose reward to the audible click of the solenoid valve. During these habituation sessions, widefield imaging was performed with the same passive stimuli used during training days to measure naïve activity and acclimate the mice to imaging. A sparse noise stimulus was presented on the last day of habituation for retinotopic mapping.

### Behavioral training

Mice were motivated to perform the task through citric acid-based water restriction^71^. Once mice had recovered from the headplate surgery, citric acid was added to the water in their home cage up to a concentration of 5%.

The behavior setup was used in part directly, and in part slightly modified, from the International Brain Laboratory design^72^. During training and recording, mice were head-fixed in the center of three screens, with all screens being gray throughout recording. Mice rested their forepaws on a wheel which could be turned leftwards (counterclockwise) or rightwards (clockwise).

The task involved turning the wheel when a visual stimulus was presented, which was rewarded with a drop of sucrose water. This task was previously described^34^, but was re-programmed for the current study using Bonsai^73^. Each trial began with a fixed inter-trial interval, which was followed by a quiescence timer that was reset by wheel movement. Mice were trained in the first one or two days with inter-trial intervals of 1-3 s and quiescence times of 0.5-1 s, and then timings were increased on subsequent days to inter-trial intervals of 4-7 s and quiescence times of 0.5-2 s. Both inter-trial intervals and quiescence times were randomly drawn on each trial. Once the quiescence timer had successfully elapsed, a stimulus appeared on the right-hand side of the mouse at 90° azimuth. The stimulus was circle spanning 20°, and contained a static square vertical grating with a spatial frequency of 0.1 cycles/degree at 100% contrast. The phase of the grating was randomly drawn on each trial. From stimulus onset, the position of the stimulus was yoked to the wheel, such that leftward movements brought the stimulus leftward on the screen towards the center, while rightward movements moved the stimulus rightward on the screen to the periphery. If the stimulus reached the center, the mouse received a 6 μL drop of 10% sucrose water. If the stimulus was moved rightward off of the screen, the trial would end.

### Associative behavior statistic

To identify sessions where mice turned the wheel in response to the stimulus, and therefore exhibited associative behavior, we used a previously described conditional randomization statistic^34,38^. Briefly, the null hypothesis is that mice turned the wheel randomly with respect to the stimulus onset. For each session, we generated a null distribution of reaction times, with reaction times being the duration from stimulus onset to first movement. The null reaction times were made using all valid stimulus presentation times for each trial, given the available set of quiescence times. In other words, for each trial, we collected all times the stimulus might have been displayed given the quiescence time options that would have resulted in the same first wheel movement. To build the null distribution of reaction times, we sampled one possible stimulus time per trial and averaged null reaction times across trials, and repeated this process 10,000 times to build a null distribution of average reaction times. We compared our measured mean reaction time to this null distribution, and rejected the null hypothesis if the measured value was at p < 0.05 in the distribution. Rejecting the null hypothesis in this case implied that the wheel turns were influenced by the stimulus, rather than the wheel being turned randomly, which is what we term “associative behavior”.

### Widefield imaging and processing

Widefield imaging was performed with an sCMOS camera (ORCA-Flash4.0 v3, Hamamatsu), mounted on a mesoscope (THT Mesoscope, SciMedia). Camera control and data collection was done by interfacing camera software (Hamamatsu) with custom MATLAB-based software. Images were binned in 2x2 pixels and collected at 70 Hz. Illumination was generated by LEDs (OptoLED, Cairn) which were blue (470 nm) or violet (405 nm), with color alternating by frame to yield a 35 Hz framerate for each color. In order to prevent light from the screens from reaching the camera, the screen was flickered out-of-phase with camera exposure. The speed of this flicker meant the screens appeared slightly darker, and flickering was not or only barely visible to the human eye.

Widefield images were compressed using singular value decomposition (SVD). SVD approximated the raw fluorescence *F* as *F* = *USV*^*T*^, where *U* is a matrix of spatial components sized *pixels x components*, and *SV*^*T*^ (below shortened to *V*) is a matrix of temporal components sized *components x time points*. The first 2000 components were saved for further processing.

Widefield data was corrected for hemodynamic effects, deconvolved to more closely reflect spiking activity, and normalized to baseline. Hemodynamic correction was performed through a combination of blue illumination, which reported activity-dependent GCaMP fluorescence, and violet illumination, which reported activity-independent GCaMP fluorescence. The process of hemodynamic correction has been described previously^13^, and involved subtracting the scaled violet signal from the blue signal. Fluorescence was then normalized to baseline and expressed as (*F* −*F*_0_)/(*F*_0_ + *s*), where *F*_0_ was the average fluorescence across the recording, and *s* was a softening factor as the median fluorescence across the average image.

Finally, deconvolution was then performed with a kernel that was previously generated to best approximate local multiunit spiking from fluorescence^13^. Applying this deconvolution kernel is very similar to taking the derivative of the fluorescence signal. Fluorescence values are labeled as dΔF/F_0_ to represent “deconvolved”, and values are more closely related to rate-of-change than total fluorescence.

Widefield data was combined across animals after first aligning images using retinotopic mapping as described in the “widefield alignment” section below. After this spatial alignment, each widefield recording still had its own set of spatial components *U*_*recording*_ corresponding to the temporal components *V*_*recording*_. In order to analyze data across recordings and animals, we performed a change of basis onto a master set of spatial components *U*_*master*_, where 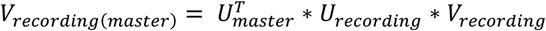. Temporal components *V*_*recording*(*master*)_ across all recordings could then be reconstructed into pixels using the same set of spatial components *U*_*master*_, which allowed us to combine temporal components across recordings for analysis.

### Widefield alignment

Widefield images were spatially aligned both within and across mice. To align days within mice, the average image from each day was aligned by rigid transformation to the first day of recording. To align across mice, we first created retinotopic maps of visual field sign using sparse noise mapping, which highlighted visual regions^74^. We then aligned the visual field sign map for each mouse to a master visual field sign map by similarity transformation.

Cortical areas from the Allen Mouse Brain Common Coordinate Framework (CCF v3, Allen Institute for Brain Science)^75^ were overlaid on widefield images by first assigning values to CCF visual regions based on their expected visual field sign^74^, then aligning that onto the master visual field sign map by similarity transformation.

### Electrophysiology recordings

Electrophysiological recordings were performed with Neuropixels Phase 3A probes and 1.0 probes (IMEC). Probes were positioned with a micromanipulator (MPM, New Scale Technologies), and probe position in the brain was approximated in real-time with integration with Neuropixels Trajectory Explorer^76^.

Electrophysiological data was collected with Open Ephys^77^.

Before each recording, after mice were head-fixed, Kwik-Cast was removed, and the craniotomy was covered with Dura-Gel (Cambridge NeuroTech). At the end of each recording, the craniotomy and Dura-Gel was covered with Kwik-Cast.

Probes were coated in dye (DiI, ThermoFisher Scientific) before every recording, and were angled at 45° or 50° down from horizontal and 90° leftward from the posterior-anterior axis, to approach the craniotomy from the right-hand side. They were inserted into the brain at approximately 0.6 mm AP and 0.7 mm ML from bregma to a depth of about 5 mm, which took the probe across the midline to diagonally enter the left striatum.

After recording, probes were cleaned by soaking in 1% Tergazyme and/or isopropyl alcohol. In cases where residual Dura-Gel stuck to the probe, the probe was soaked in DOWSIL DS-2025 cleaning solvent.

Electrophysiology data was processed by first performing common average referencing. This was done by first taking the median across simultaneously sampled channels, scaling that median to each channel by regression, then subtracting the scaled median from each channel. This removed most of the electrical noise and imaging-induced artifacts. Spikes were then sorted with Kilosort4^78^, and units representing noise were identified using Bombcell^79^ and removed from further analysis.

The borders of the striatum were defined for each recording based on electrophysiological landmarks. The start of the striatum was identified after a gap in units corresponding to the ventricle, and the end of the striatum, when present, was identified as a sudden shift in multiunit correlation near the bottom of the probe as it entered the endopiriform cortex or claustrum. Occasionally, issues with the probe caused less than 50 units to be detected in the striatum, and these recordings were discarded. Also, on occasion, the probe passed through the external globus pallidus (GPe) instead of the striatum. These recordings were clearly evident from high firing rates, and were also excluded.

After experiments were completed, mice were perfused with 4% formaldehyde, brains were fixed overnight at 4°C, then transferred to PB for at least 24h. The brain was then sectioned and imaged with a fluorescence microscope (SMZ25, Nikon).

### Corticostriatal maps and clustering

Functional mapping from cortical areas to striatal domains was done through ridge regression as previously described^13^. Striatal activity was binned into multiunit activity in 200 µm segments across each recording, the first and last 60 seconds of data were excluded, and spike rates were z-scored. Cortical activity, being the first 200 temporal components (*V*), was then regressed to striatal activity. Ridge regression was performed with a regularization value λ = 10, and the design matrix included temporal lags from -200 to 200 ms. The cortical map for each striatal segment was the maximum kernel value across time lags.

Corticostriatal map clustering was performed using k-means across maps from all striatal segments across all recordings. K-means was set to produce 3 clusters, which empirically captured the variability seen across maps. Note that the corticostriatal gradient is continuous rather organized into discrete domains, but this particular trajectory through the striatum is well-characterized by being grouped into 3 domains^13^. Initial clusters for k-means were seeded by averaging maps across thirds of each recording, then averaging across recordings, producing average maps for the upper, middle, and lower segments of the probe in the striatum. Each 200 µm segment from all striatal recordings was then assigned into a domain based on the k-means clustering result.

### Cortical regions of interest corresponding to striatal domains

Cortical regions of interest (**Supplementary Fig. 3C, yellow outlines**) were generated from the average corticostriatal map within each cluster. The average map from each domain was gaussian filtered with a standard deviation of 10 pixels, and a region of interest (ROI) was defined where the filtered image value was greater than 0.75 times the maximum value. These ROIs represented the cortical regions directly upstream of each striatal domain.

Fluorescence was then extracted for each trial in these ROIs. Fluorescence was already amplitude-normalized from the widefield preprocessing, and baseline-subtraction was performed for each trial using the average fluorescence in a 500 ms window before stimulus onset.

### Striatal electrophysiology analysis

Striatal cell types were classified using parameters extracted from Bombcell^79^. Narrow and wide waveforms were defined from a peak-to-trough threshold of 400 µs, and bursty and regular-spiking units were defined from a post-spike suppression threshold of 40 ms. Medium spiny neurons (MSNs) were defined through wide waveforms and regular spiking, fast-spiking interneurons (FSIs) were defined through narrow waveforms and regular spiking, and tonically active neurons (TANs) were defined through bursty spiking (**Supplementary Fig. 5**). Narrow waveforms with two peaks were classified as putative axons and not included in analysis.

For analyses examining multiunit activity in the striatum (**Fig. 1-3, Supplementary Fig. 4**), only spikes from putative MSNs were included, and spikes from all MSNs within a given domain were included (schematized in **Supplementary Fig. 3E-F**). Multiunit firing rate (FR) was normalized as ΔFR/FR_0_ = (*FR*−*FR*_0_)/(*FR*_0_ + *s*), where *FR*_0_ was the average firing rate 500 ms before stimulus onset on each trial, and *s* was a softening factor of 10 spikes/s. Multiunit firing rate was then smoothed with a causal half-gaussian filter with a width of 100 ms. Single unit data (**Fig. 4**) was normalized and smoothed in the same way, but with a softening factor *s* of 1 spike/s.

Units were identified as being significantly responsive to passive stimulus presentations (**Fig. 4**) through a shuffle test. Spikes were counted in an 50-150 ms window before and after the stimulus. The average difference between pre- and post-stimulus spiking was compared to shuffling the pre- and post-stimulus spike counts within trials 10,000 times. If the firing rate increased after stimulus onset at p > 0.99 relative to the shuffle distribution, the unit was classified as being responsive to that stimulus. This was done separately for each of the three passively presented stimuli on the left, center, and right-hand screens.

## Supplementary Figures

**Supplementary Figure 1.**
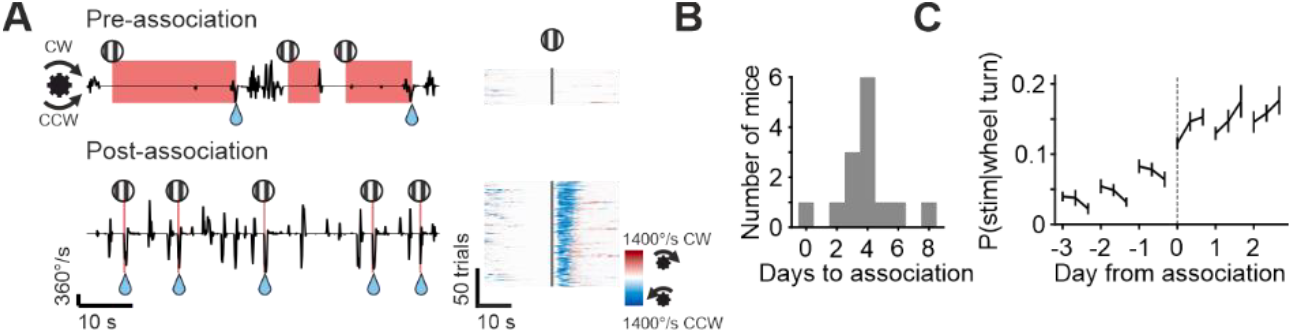
Visuomotor associative behavior. (A) Example wheel velocity in a session before and after learning the association. Left, snippet of wheel velocity with labels for stimulus presence (stimulus symbol, red shading) and reward delivery (water drop symbol), positive values are rightward/clockwise turns and negative values are leftward/counterclockwise turns; right, wheel velocity aligned to all stimulus onsets within the session. (B) Histogram of number of mice which exhibited associative behavior on each day of training. There was one mouse which did not learn the association throughout training, which is plotted at “0 days to learn”. (C) The fraction of all wheel turns during the task which were preceded by a stimulus onset within 400 ms. Sessions are aligned relative to the first association day within each mouse, and trials are split into thirds within each session. Curves and error bars are mean ± SEM across mice. Note the within-day downward trend before association suggesting fewer stimulus-movement overlaps by chance, and the within-day upward trend after association suggesting more efficiently executed movement to the stimulus.

**Supplementary Figure 2.**
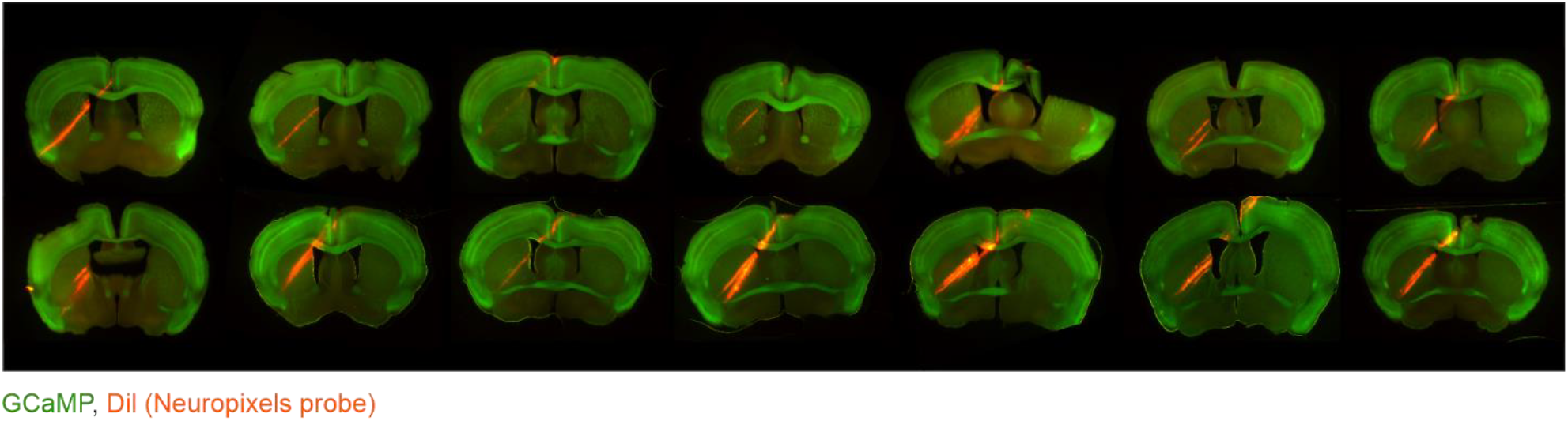
Striatal electrophysiology location. One example histology slice from each mouse, showing GCaMP expression (green) and probe tracts through the striatum (DiI, orange).

**Supplementary Figure 3.**
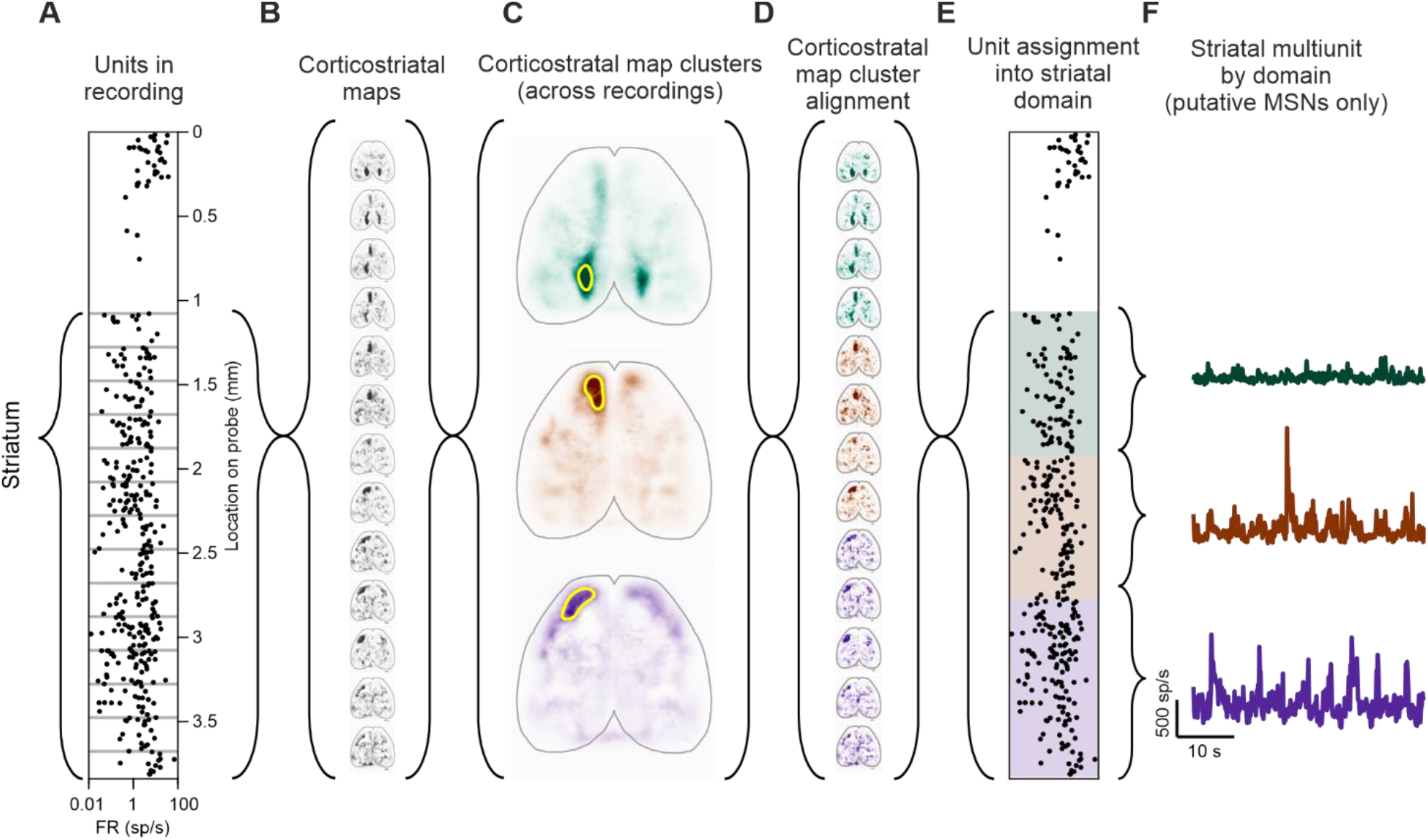
Defining striatal domains by correlated cortical regions. (A) Example units recorded in one session, plotted as position on probe against spike rate across the session. The location of the striatum is noted, which begins after the gap in units on the part of the probe in the ventricle. Gray lines denote 200 µm segments used for assigning striatal domain. (B) Example corticostriatal maps for each segment in (A), which represent kernels from regressing widefield fluorescence to striatal multiunit activity within each segment. (C) Mean clustered corticostriatal maps from all striatal segments across all recordings, corresponding to domains within the striatum. Regions-of-interest are plotted for each map (yellow) which were used to extract cortical activity associated with each striatal domain (e.g. plotted in Fig. 2A, green). (D) Corticostriatal maps in (B), colored according to their assigned domain from the clustering in (C). (E) Units in (A), with each striatal segment colored according to its assigned domain from the clustering in (C). (F) Example multiunit activity by striatal domain from recording in (E), created by combining the spikes of all medium spiny neurons within each domain (e.g. plotted in Fig. 2A, black).

**Supplementary Figure 4.**
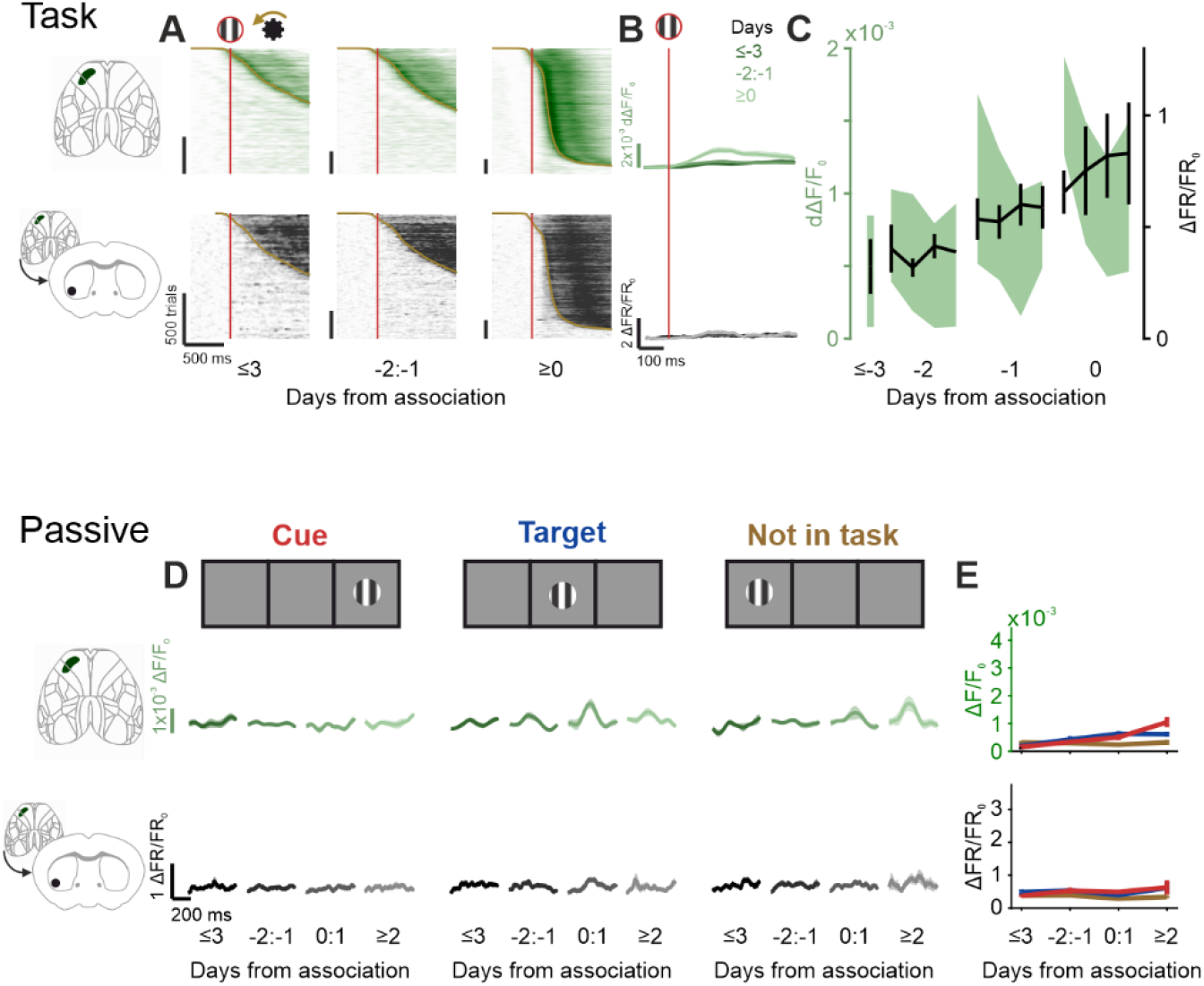
Orofacial-recipient striatum does not exhibit visual responses. (A-C) Data as in Fig. 2A-C, for orofacial cortex and orofacial-recipient striatum. (D-E) Data as in Fig. 3B, E, H, J, for orofacial cortex and orofacial-recipient striatum.

**Supplementary Figure 5.**
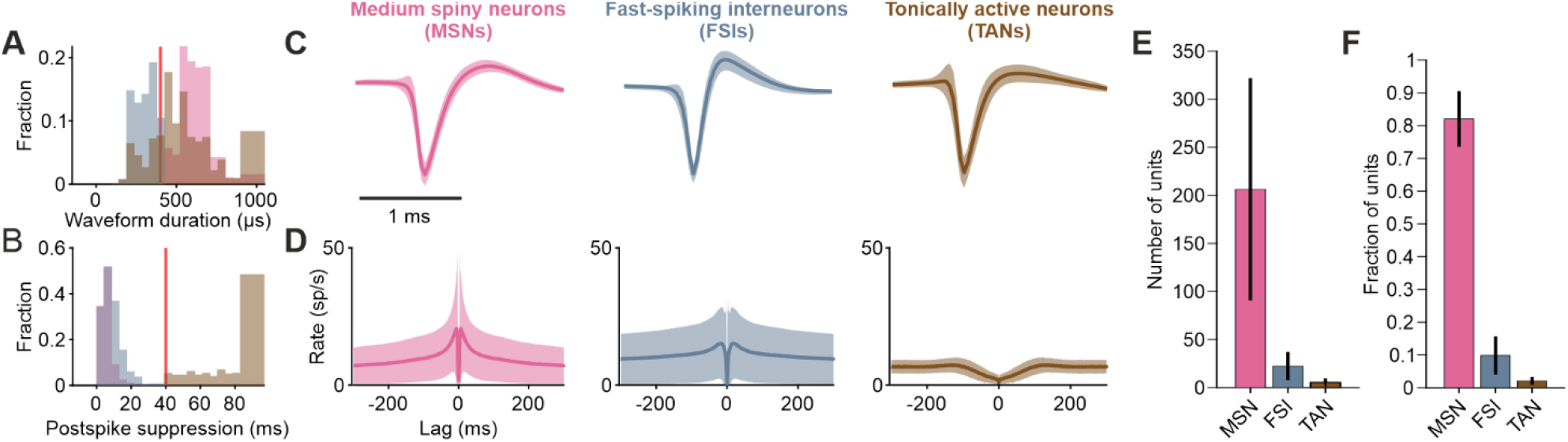
Striatal cell types can be identified by electrophysiological properties. (A) Histogram of waveform durations by striatal cell type, normalized to proportion within each cell type. Waveform duration is defined from trough to peak. Red line denotes threshold between narrow (under) and wide (over) waveforms. Pink, medium spiny neurons (MSNs); gray, fast-spiking interneurons (FSIs); brown, tonically active neurons (TANs). (B) Histogram of post-spike suppression times, normalized and colored as in (A). Post-spike suppression is defined as the time for the autocorrelation to reach baseline after a spike, with baseline being defined as the average autocorrelation value between lags of 100-300 ms. Red line denotes threshold between bursty (under) and regular-spiking (over) types. (C) Average waveforms within each striatal cell type. Waveforms of each unit are normalized to maximum absolute amplitude within each unit. Lines and shading are mean ± SD. (D) Average autocorrelation within each striatal cell type. Lines and shading are mean ± SD. (E) Average number of units classified as each striatal cell type per recording, bars and error bars are mean ± SD across recordings. (F) Average fraction of striatal units classified as each cell type per recording, error bars are mean ± SD across recordings.

